# A First Ultrastructural and Immuno-Labelling Investigation of Western Red Cedar Leaves

**DOI:** 10.1101/2023.01.28.526058

**Authors:** Brent E. Gowen, David Noshad

## Abstract

A first morphological analysis of the leaves of *Thuja plicata* (Western Red Cedar, WRC), a commercially valuable tree species, is presented. The ultrastructural and immuno-labelling analyses were only possible after applying modifications to standard transmission electron microscopy (TEM) preparatory methodology to fully embed cedar leaves. We demonstrate an application of the technique after leaf exposure to the fungi *<u>Didymascella thujina</u>* (cedar leaf blight, CLB). The leaves resulting defence response was visualized to two plant defence proteins, β-1,3 Glucanase and Chitinase. The TEM methodology modifications presented may generally apply to other plant species resistant to morphological analyses due to their thick leaf cuticles. The technique requires no more equipment than found in any basic Transmission Electron Microscopy Facility.

## Introduction

Western Red Cedar (*Thuja plicata*) is a commercially valuable tree species native to the Pacific Northwest of North America. Its usage includes lumber, shingles, and ornamental foliage. Many diseases can affect WRC via infection through the root system. However, *<u>Didymascella thujina</u>* can invade via the leaves, causing Cedar Leaf Blight. The disease appears most commonly on young seedlings and the lower branches of older trees. Young seedlings and saplings sustain the most damage where stem or branch death may occur. Disease on trees older than 4-5 years can retard growth. Disease levels are highest in dense stands where high humidity levels are high, such as in plant nurseries. The disease can be a serious problem for developing young plants (Kope and Sutherland 1994).

Considering its economic importance, it is surprising that there are no ultrastructural studies of WRC and how environmental or genetic factors can affect its normal growth and susceptibility to diseases. We have been unable to find any Transmission Electron micrographs of WRC leaves under either normal, or disease conditions.

The fixation and plastic embedding of plant material for TEM are difficult due to their inherent characteristics such as thick cellulosic or lignified cell walls, waxy substances within hydrophobic cuticles, large amounts of gases in the intercellular spaces, and the presence of large water-filled vacuoles. Special modifications of the protocols generally used for standard TEM embedding are often required (Kuo, 2007). The plant cuticle is an extracellular hydrophobic layer that covers the aerial epidermis of all land plants, protecting against desiccation and external environmental stresses (Yeats and Rose, 2013). It consists of lipid and hydrocarbon polymers impregnated with wax and is synthesized exclusively by the epidermal cells (Kolattukudy 1996). The cuticle functions in defence for the plants, forming a physical barrier that resists penetration by virus particles, bacterial cells, and the spores and growing fungal filaments (Freeman 2002) and (Noshad 2023).

Plants produce a number of defense chemicals that aid in their defense against invasive pathogens. β -1,3 glucanase is an enzyme that breaks down β-1,3-glucans found in the cell walls of fungi. Chitinases are hydrolytic enzymes that break down glycosidic bonds in chitin. (Jollès P. and Muzzarelli R.A. 1999). As chitin is a component of the cell walls of fungi, chitinases are generally found in fungi to reshape their own chitin (Sámi et al. 2001). β-Glucans are a group of β-D glucose polysaccharides naturally occurring in the cell walls of fungi.

We report on the morphological methodologies required to investigate ultra-structurally normal and diseased WRC leaves and enable accurate plant defence protein immuno-localizations.

## Methods

### Cedar Leaf Samples and Fixation

Young seedling Western Red Cedar trees were exposed to *Didymascella thujina*. Visually infected leaves were cut from their branches and placed into 0.1 M buffered glutaraldehyde and formaldehyde (both at 3%) (Hayat 1989). Samples were also collected from age-matched control, uninfected trees. The samples were fixed in specimen vials for three hours at room temperature under a vacuum and stored overnight in the fridge. While many sample pieces had sunk in the fixative, many remained floating. After warming to room temperature, all the floating pieces were removed the next day. Only the pieces that had sunk into the fixative were processed further. Table 1 summarizes many fixation and embedding procedures attempted to embed WRC leaves entirely for morphological investigation based on principles from Hayat (1989), Skepper (2000) and Newman and Hobot (1999).

**Table 1.**
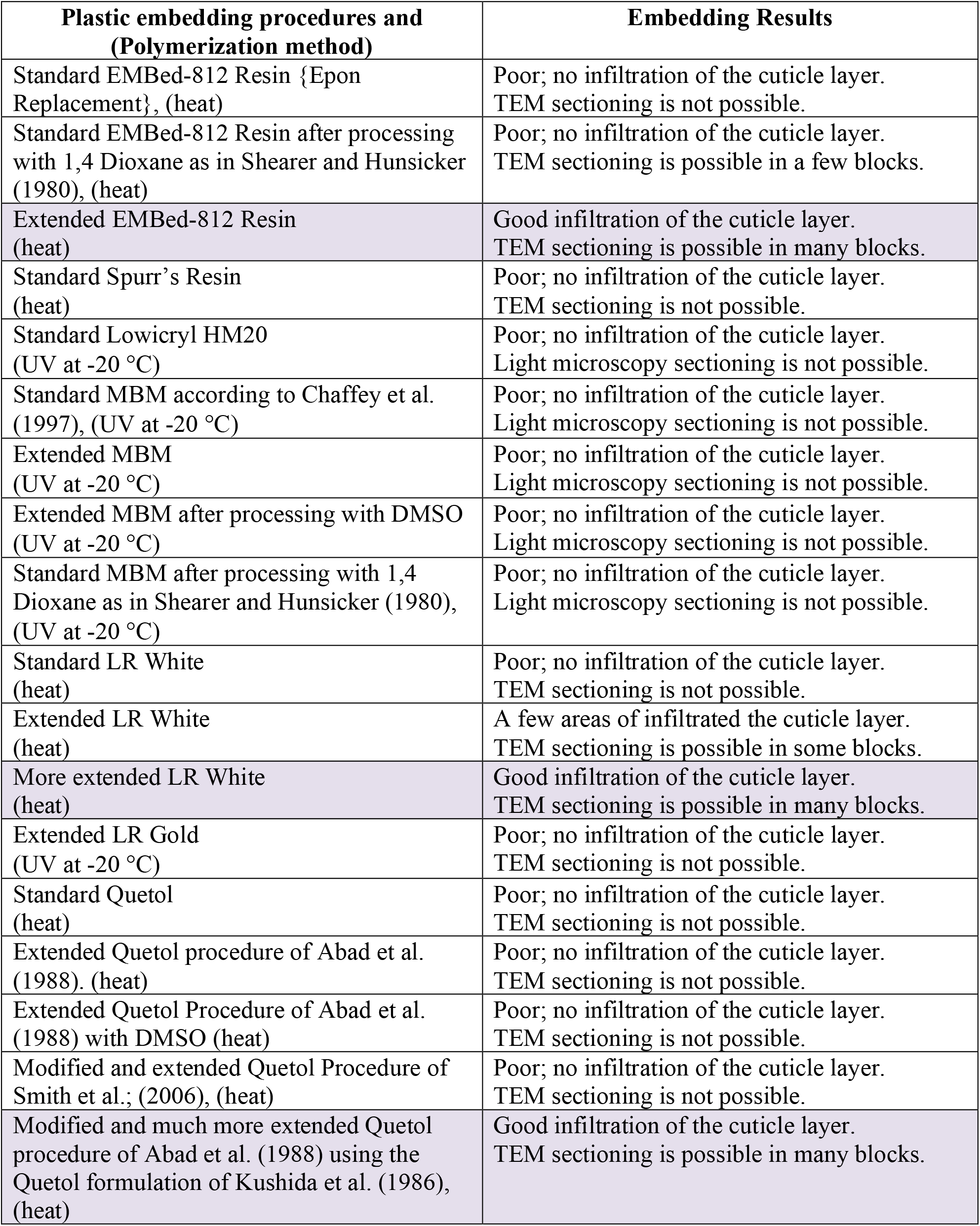
All procedures table. Shaded procedures include procedures and images that are presented within this paper.

| **Plastic embedding procedures and (Polymerization method)** | **Embedding Results** |
| --- | --- |
| Standard EMBed-812 Resin {Epon Replacement}, (heat) | Poor; no infiltration of the cuticle layer. TEM sectioning is not possible. |
| Standard EMBed-812 Resin after processing with 1,4 Dioxane as in Shearer and Hunsicker (1980), (heat) | Poor; no infiltration of the cuticle layer. TEM sectioning is possible in a few blocks. |
| Extended EMBed-812 Resin (heat) | Good infiltration of the cuticle layer. TEM sectioning is possible in many blocks. |
| Standard Spurr's Resin (heat) | Poor; no infiltration of the cuticle layer. TEM sectioning is not possible. |
| Standard Lowicryl HM20 (UV at -20 °C) | Poor; no infiltration of the cuticle layer. Light microscopy sectioning is not possible. |
| Standard MBM according to Chaffey et al. (1997), (UV at -20 °C) | Poor; no infiltration of the cuticle layer. Light microscopy sectioning is not possible. |
| Extended MBM (UV at -20 °C) | Poor; no infiltration of the cuticle layer. Light microscopy sectioning is not possible. |
| Extended MBM after processing with DMSO (UV at -20 °C) | Poor; no infiltration of the cuticle layer. Light microscopy sectioning is not possible. |
| Standard MBM after processing with 1,4 Dioxane as in Shearer and Hunsicker (1980), (UV at -20 °C) | Poor; no infiltration of the cuticle layer. Light microscopy sectioning is not possible. |
| Standard LR White (heat) | Poor; no infiltration of the cuticle layer. TEM sectioning is not possible. |
| Extended LR White (heat) | A few areas of infiltrated the cuticle layer. TEM sectioning is possible in some blocks. |
| More extended LR White (heat) | Good infiltration of the cuticle layer. TEM sectioning is possible in many blocks. |
| Extended LR Gold (UV at -20 °C) | Poor; no infiltration of the cuticle layer. TEM sectioning is not possible. |
| Standard Quetol (heat) | Poor; no infiltration of the cuticle layer. TEM sectioning is not possible. |
| Extended Quetol procedure of Abad et al. (1988). (heat) | Poor; no infiltration of the cuticle layer. TEM sectioning is not possible. |
| Extended Quetol Procedure of Abad et al. (1988) with DMSO (heat) | Poor; no infiltration of the cuticle layer. TEM sectioning is not possible. |
| Modified and extended Quetol Procedure of Smith et al.; (2006), (heat) | Poor; no infiltration of the cuticle layer. TEM sectioning is not possible. |
| Modified and much more extended Quetol procedure of Abad et al. (1988) using the Quetol formulation of Kushida et al. (1986), (heat) | Good infiltration of the cuticle layer. TEM sectioning is possible in many blocks. |

### EMBed-812 Resin (Epon replacement) Processing

After washing in buffer three times, the samples were osmicated with 1.0% osmium tetroxide in buffer for one hour. After washing in buffer three times and a graded ethanol dehydration (all steps for 20 minutes and at room temperature on a rotor), the transition fluid of 100% propylene oxide was used twice for 20 minutes each time. The samples were left on the rotor in propylene oxide/812 Resin with no catalyst (3:1) for one day, propylene oxide/812 Resin with no catalyst (1:1) for one day, propylene oxide/812 Resin with no catalyst (1:3) for five days, pure 812 Resin with no catalyst for two days, new pure 812 Resin with no catalyst for four days, new 812 Resin with catalyst for one day, new 812 Resin with catalyst for two days, and then the samples were embedded into new 812 Resin in moulds and polymerized at 60° C for two days.

### Quetol Embedding Procedure

Modified from Abad, Cease, and Blanchette (1988) and using the modified Quetol recipe from Kushida et al. (1986); Quetol 651: 28 g, ERL 4206 (an old bottle of the original component within the UVic EM Lab): 6 g, NSA: 66 g and DMP-30: 1.5–2.0 g. After washing in buffer three times, the samples were osmicated with 1.0% osmium tetroxide in buffer for one hour. After washing in water three times and a graded ethanol dehydration (all steps for 20 minutes and at room temperature on a rotor), the samples were left on the rotor in 75% Quetol 651: 25% for five days, 100% Quetol 651 for two days, new 100% Quetol for three days, Kushida Quetol formulation without DMP-30 for four days, Kushida Quetol formulation with DMP-30 for five days, and then samples into moulds and polymerized the plastic for two days at 60°C.

### LR White Processing

After washing in buffer three times and a graded ethanol dehydration (all steps for 20 minutes and at room temperature on a rotor), the samples were left in 100% ethanol/LR White monomer (1:1) for five days on the rotor. This was followed by 100% ethanol/LR White monomer (1:3) for three days, pure LR White monomer for one day and new pure LR White monomer for 4 hours and then 12 days in the fridge. After this time, the sample was warmed and placed into new LR White monomer for five days on a rotor. Individual leaf pieces were then placed into individually labelled gelatin capsules. The capsules were covered by their second half and put into a polymerization oven at 60°C for one day.

### Chemicals Used

#### Primaries

Antibodies to β-1,3 Glucanase and Chitinase were purchased from Agrisera AB Box 57, SE-911 21 Vännäs, SWEDEN. Both primaries are polyclonal rabbit IgGs and diluted at 1:600. An antibody to (1,3)-β-Glucan (monoclonal mouse IgG) was purchased from Biosupplies Australia Pty Ltd, PO Box 187, La Trobe University, Bundoora, VIC 3083, Australia) and diluted at 1:300.

#### Secondaries

Fluorophore-conjugated secondary antibodies (AlexaFluor 488 and AlexaFluor 568) were purchased from Invitrogen/Thermo Fisher Scientific, 168 Third Avenue, Waltham, MA, USA 02451. Both were diluted at 1:100.

The nuclear fluorescent stain was DAPI (Invitrogen/Thermo Fisher Scientific D1306) 10.9 mM aliquot diluted 1:50 in buffer.

Colloidal gold conjugated secondary antibodies to mouse and rabbit primaries were purchased from Jackson ImmunoResearch Inc 872 W. Baltimore Pike, West Grove, PA USA 19390). 12^nm^ Gold-AffinitiPure Goat anti-rabbit IgG (H+L), Code Number 115-205-144 and 6^nm^ Gold-AffinitiPure Goat anti-mouse IgG (H+L), Code Number 115-195-146.

All TEM-associated chemicals and embedding plastics were purchased from Electron Microscopy Sciences (EMS), P.O. Box 550, 1560 Industry Road, Hatfield, PA 19440, USA.

### Light Microscopy Immunolabeling of LR White Sections

The immunolabelling procedures were modified from Liljeroth, Marttila and von Bothmer (2005). Sections from LR White blocks were cut at 1.0^μm^ thickness with a Histo diamond knife (Diatome Ltd., Helmstrasse 1 2560 Nidau, Switzerland) and placed onto Superfrost Plus slides (Fisher Scientific). Groups of sections were isolated using a PAP pen (Sigma-Aldrich Canada) and then blocked using PBS with 1% ovalbumin for 20 minutes. The primaries, secondaries and DAPI were diluted in the blocking solution. After labelling, the slides were mounted with Fluoromount G (EMS) and sealed with nail polish.

A Leica DM LB2 compound light microscope with a UV Fluorescence EBQ 100 Lamp Control Unit and DFC425 digital camera controlled with V3.5.0 Leica Application Suite software were used to collect images. Images were compiled using Adobe Photoshop, version 12.1 (Adobe Systems Inc., San Jose, CA, USA).

### TEM Immunolabelling of LR White Sections

The TEM immunolabelling procedures followed were from von Schalburg et al. (2018). Silver/gold interference-coloured sections were cut of all blocks. The sections were placed onto 100 mesh copper grids containing carbon-coated Formvar™ films. The grids were blocked using PBS with 1% ovalbumin for ten minutes in a humid chamber. The primaries and secondaries were diluted in the blocking solution. After labelling, the sections were stained in 5% uranyl acetate in 50% ethanol for ten minutes and 5% lead citrate for two minutes. Sections were viewed on a JEOL 1011 Transmission Electron Microscope operating at 80 kV, and images were captured using a Gatan ES 100W Erlangshen CCD camera. No immunolabelling occurred when the primaries were eliminated from the procedure (data not shown).

## Results

For standard TEM processing, small pieces of a sample are required to allow for the complete and timely infiltration of fixatives, dehydrators, and embedding plastic. The recommended size beginning with immersion fixation for plants is 0.5-1.0 mm^3^ in size (Hayat, 1989). Even when cutting our WRC samples this small, these pieces did not fully embed using standard TEM processing methods when the cuticle was present. With this material missing within embedded tissue, obtaining good 1.0 um survey sections or ultrathin sections (75-85 nm thin) was not possible. As this is the site of fungal infection of the leaves, this cannot be investigated after “standard” TEM processing of WRC leaves.

**Table 1** summarizes many fixation and embedding procedures tried to embed WRC leaves completely for morphological investigation. We were limited against possible other techniques, such as microwave-assisted processing (Clode 2015), Tokuyasu’s cryo-sectioning method (1978) and vitrification (high-pressure freezing) followed by freeze substitution (McDonald 2014), as we have access to only a basic Electron Microscope Facility. The alternative techniques require expensive equipment and training beyond our, and many, EM facilities.

**Figure 1**. shows a fully embedded cuticle from the extended Quetol procedure.

**Figure 1.**
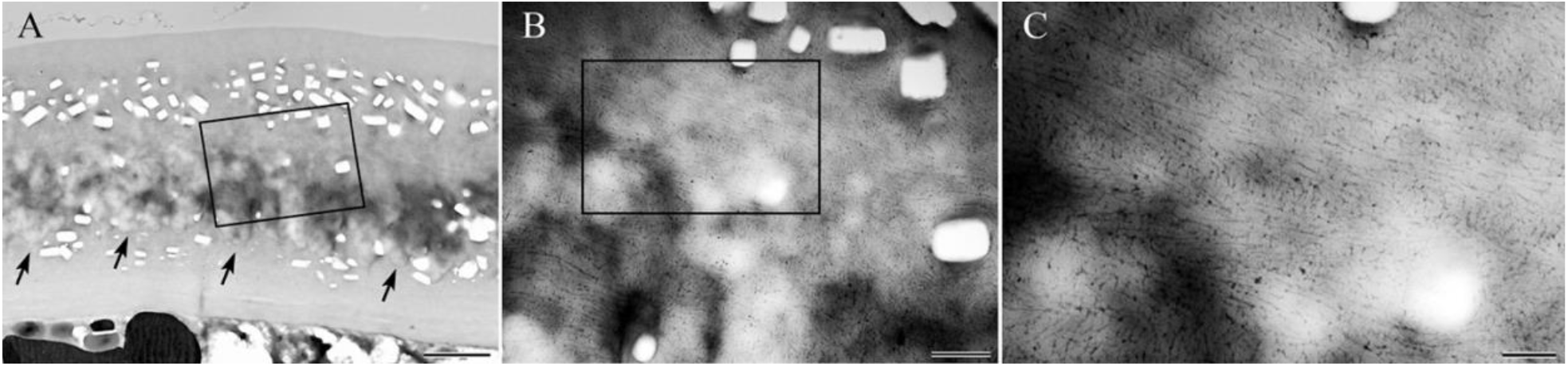
Extended Quetol embedded cedar leaf. All pictures are from the same area of a section. A = low magnification image shows the thick wax layer within the leaf’s cuticle. While the dorsal part is less distinct as to where the wax starts, the arrows point to the ventral part where the layer of wax has a distinct end within the cuticle. The boxed area indicates where part B of the figure originates. The clear crystalline areas within the cuticle are the calcium oxalate “ghosts” where the oxalate dissolved during processing. Scale bar = 2 µm. B = higher magnification of the wax layer showing the structured layer nature of the wax. The boxed area indicates where part C of the figure originates. Scale bar = 0.5 µm. C = higher magnification of the wax layer showing the fine detail arrangement of the wax particles within the cuticle. Scale bar = 200 nm.

**Figure 2**. shows a fully embedded cuticle from the extended EMBed-812 procedure.

**Figure 2.**
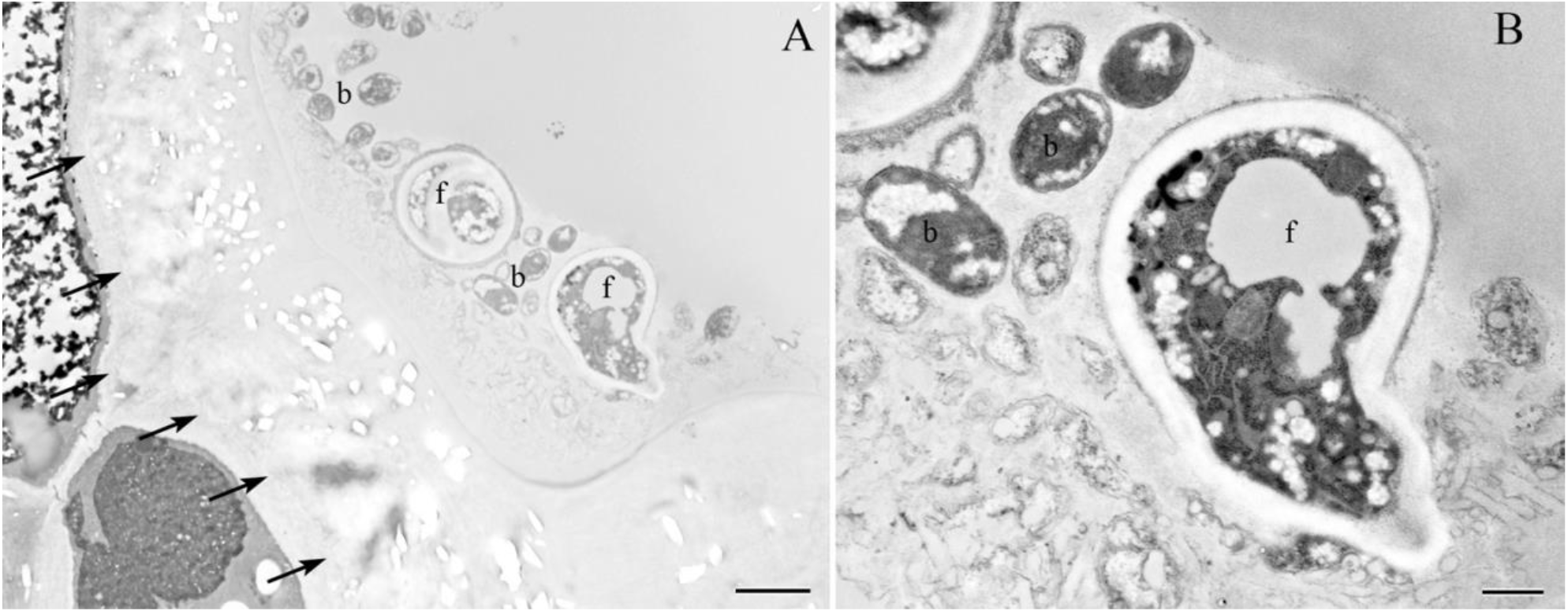
Extended EMBed-812 embedded cedar leaf. All pictures are from the same area of a section. A = low magnification image showing the thick layer of wax within the fully embedded cuticle of a leaf. As seen in the Fig 1, the ventral part of the wax is well delineated (arrows). Calcium oxalate ‘ghosts’ are present. Fungi (f) and bacteria (b) are located outside the leaf. Scale bar = 2.0 µm. B = a higher magnification closeup of A. Scale bar = 0.5 µm.

**Figure 3**. shows double immunofluorescence labelling of leaf defence proteins and fungus. There is little response by defence proteins where the fungus is present only on the outside. When the fungus is inside the leaves, there are much more defence proteins in the immediate vicinity of the fungus.

**Figure 3.**
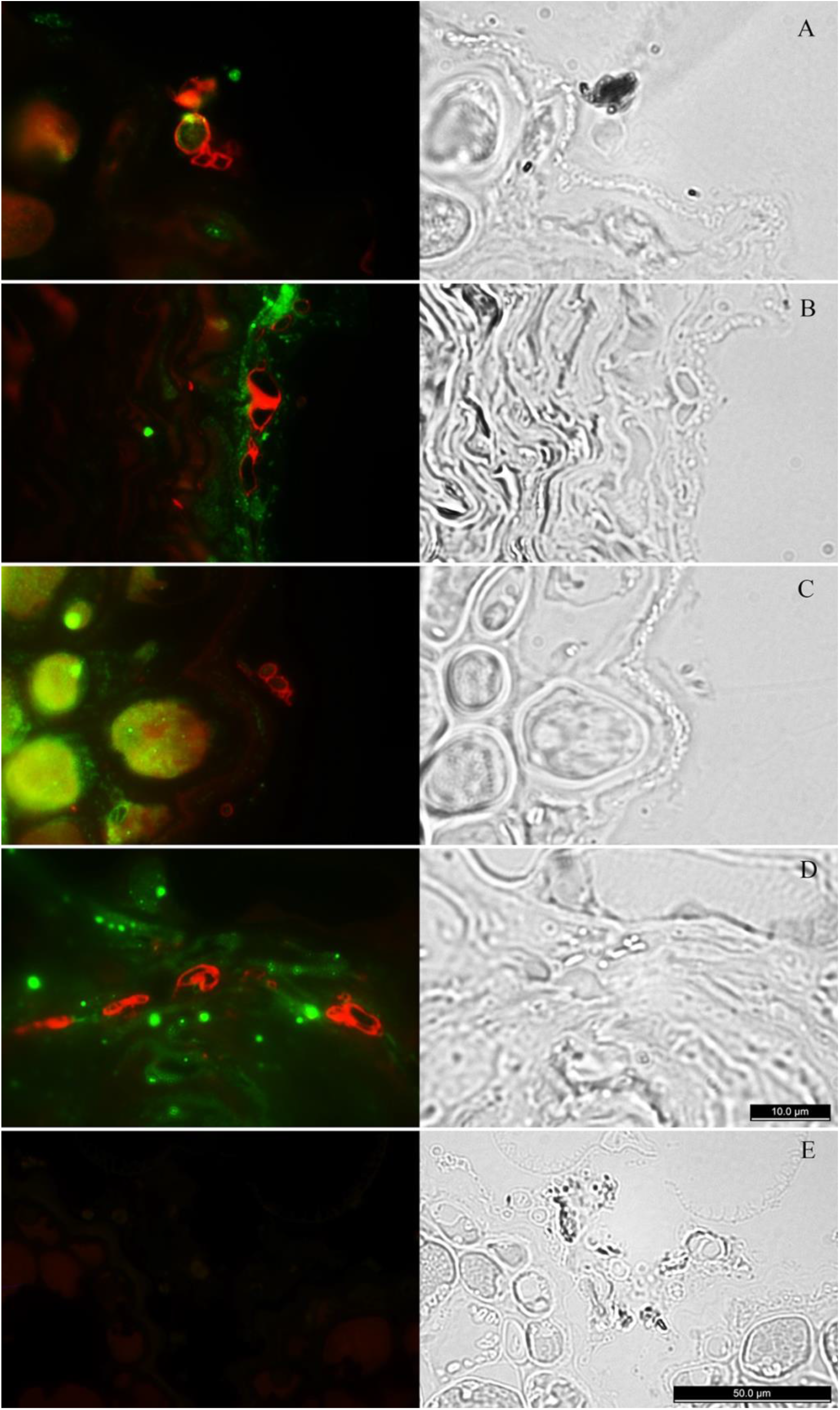
LM Immunolabelling of extended LR White embedded cedar leaf. A = chitinase (red) and B-glucan (green). Fungus on the outside of cuticle, no defence reaction to the fungus. B = chitinase (red) and B-glucan (green). Fungus on the inside of cuticle with defence reaction (green immunolabelling) to the fungus. C = gluconase (red) and B-glucan (green). Fungus on the outside of the cuticle has no defence reaction to fungus (but a possible strong response to something further within the leaf, like infection from the other side of the leaf ?). D = gluconase (red) and B-glucan (green). Fungus on the inside of the cuticle with defence response to the fungus. E = PBS controls: no primaries present in the immunolabelling run, but secondaries were. The fluorescent image is on the left side of every figure while the right side is the corresponding phase contrast image. For A) to D); scale bar = 10 µm. For E); scale bar = 50 µm.

**Figure 4**. shows double TEM immunolabelling of leaf defence proteins and fungus. The fungus (and bacteria) is within the leaf. The images show the specificity of the labels as they are specific for the fungal cell wall and defence proteins around the fungus. The invading bacteria present are unlabelled.

**Figure 4.**
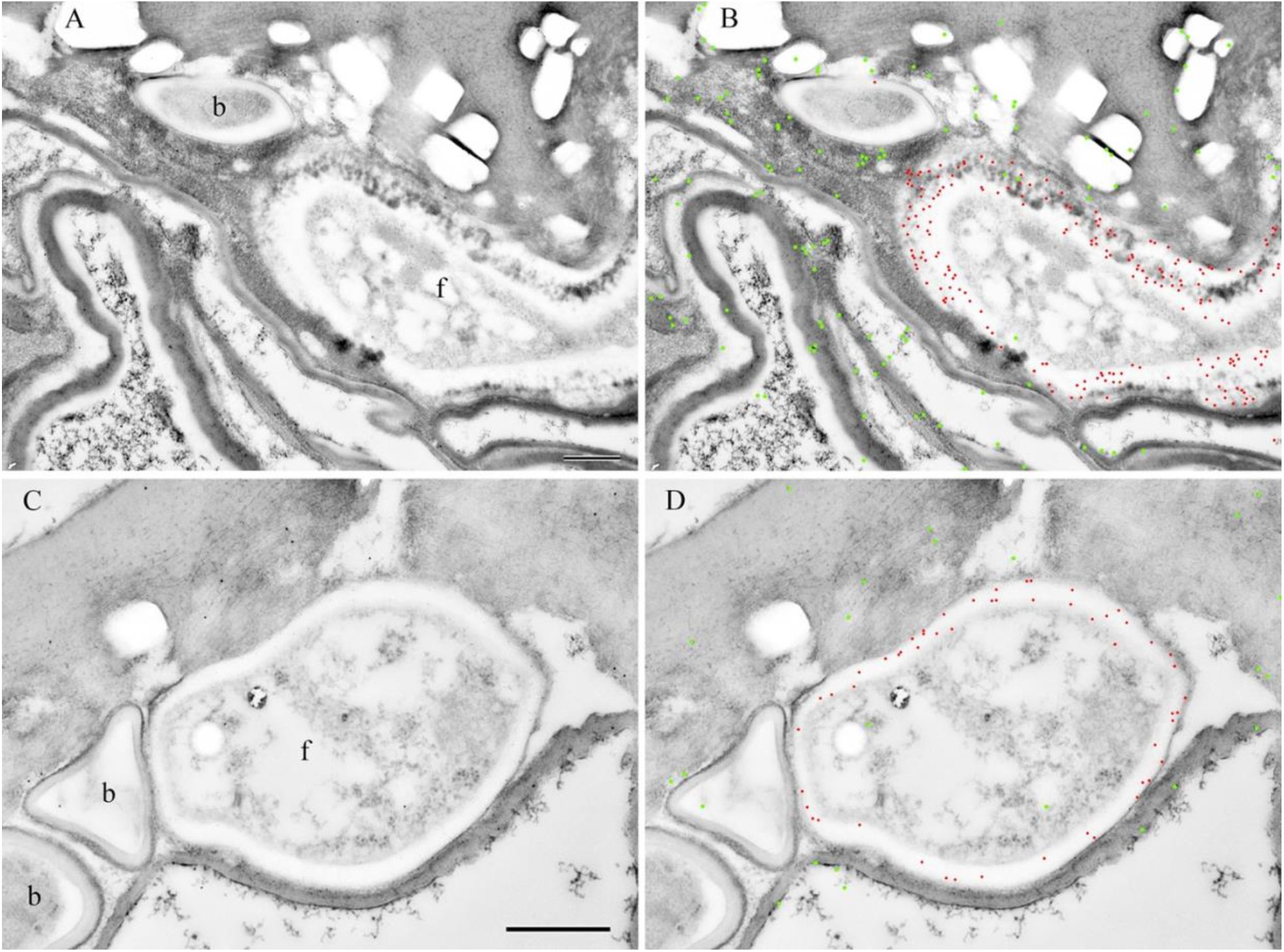
TEM Immunolabelling of extended LR White embedded cedar leaf. The fungus is within the leaf in these images. f = fungus, b = bacteria. Note that the bacteria are not labelled. The labelling is separate and precise for the fungus and defence proteins. A = {top left} LR White section double labelled for β -1,3 glucanase with a 12 nm gold secondary and for β-Glucan with a 6 nm secondary. Scale bar = 0.5 µm. B ={top right} is a false coloured version of A) where the 12 nm gold is coloured green and the 6 nm gold red (to clarify the results as 6 nm gold is not as apparent at this scale unless coloured). C = {bottom left} LR White section double labelled for Chitinase with a 12 nm gold secondary and β-Glucan with a 6 nm secondary. Scale bar = 0.5 µm. D = {bottom right} is a false coloured version of A) where the 12 nm gold is coloured green and the 6 nm gold red (to clarify the results as 6 nm gold is not as apparent at this scale unless coloured).

## Discussion

Processing biological tissue for transmission electron microscopy involves several steps, including removing the tissue from the whole organism, fixation of proteins and lipids, removal of water, and the infiltration of plastic. These steps are optimized for morphological presentation as close to the live material as possible with minimal or known artifacts (Hayat, 1989). When the processing is to be combined with immuno-labelling procedures, one must ensure that the processing steps do not alter the locations of the antigens of interest from their usual sites or alter them to the point that immunoglobulins can no longer recognize, and therefore label, them (Polak and Van Noorden 1997) and (Skepper 2000). Kuo (2007) advocated for rapid conventional chemical preparation methods (up to eight hours) for ultrastructural studies of plant cells to minimize artifact formations caused by prolonged specimen preparation generally used for embedding biological samples. Even after our extended embedding procedures, we show that antigens of interest for investigating cedar’s response to invading fungal disease agents; (a) are visualized using immunolabelling procedures at both the light and electron microscope levels using the same embedded blocks and (b) the antigens remain in their original locations.

Fungal pathogens breach plant cuticles using a combination of enzymatic degradation and mechanical rupture. The latter is often accomplished by forming a swollen appressorium structure that extends an infectious peg via turgor pressure (Deising et al., 2000). While mechanical rupture may be sufficient for cuticle penetration, particularly of thinner cuticles (Tenberge,2007), most fungal pathogens also secrete cutinases, a class of small, nonspecific esterases that hydrolyze the cutin polyester and release free cutin monomers (Longhi and Cambillau 1999). The cutin monomers released during polymeric cutin hydrolysis can act as elicitors of plant defence responses and are thus classified as damage-associated molecular patterns. At micromolar concentrations, these compounds induce the production of hydrogen peroxide and other defence responses (Schweizer et al., 1996 and Kauss et al., 1999).

While we do not show the initial penetration of the fungus into the leaf, we now have the technical ability to view how this happens with *<u>Didymascella thujina</u>* as it infects Western Red Cedar (*Thuja plicata*). In addition, we can now investigate, combined with biochemical techniques, how environmental and genetic modifications of WRC may aid in minimizing the fungal infection of this valuable commercial species.

We obtained accurate cuticle ultrastructure using the modified EMBed-812 and Quetol procedures. However, these plastics are not generally useful for immunolabelling light and transmission electron microscopy sections. The immunoglobulins do not penetrate beyond the surface of these sections. The osmication and high-temperature polymerization steps necessary for embedding into these plastics often denature many antigens of interest, reducing the chances for good immunolabelling of sections (Skepper, 2000). Embedding the WRC into LR White is a good compromise for finding our antigens of interest in light and transmission electron microscopy sections in this difficult-to-embed tissue.

The TEM methodology modifications may generally apply to other plant species resistant to morphological analyses due to their thick leaf cuticles. The technique requires no more equipment than found in any basic Transmission Electron Microscopy Facility.

## Notes

### Competing Interest Statement

The authors have declared no competing interest.

